## Supplemental material S1_Text for "Comparative analysis of shared and unique mechanisms important for diverse strains of *Pasteurella multocida* to cause systemic infection in mice"

Sequencing and assembly of the *P. multocida* strain M1404 genome

Prior to TraDIS analysis we performed whole-genome sequencing of *P. multocida* strain M1404 via Illumina and Nanopore sequencing as described in the materials and methods. The M1404 genome was assembled as previously described [1], with minor modifications. Briefly, Nanopore data were processed using High Accuracy Calling, and then further processed using Filtlong to remove reads < 1 kb in length and 5% of reads that had the lowest quality score. The processed M1404 nanopore reads were *de novo* assembled using Flye and then polished with Nanopore reads using Medaka. Illumina M1404 reads were trimmed using Trimmomatic, then used to polish the Nanopore assembly using Polypolish and then POLCA. The genome was then annotated using Prokka. The M1404 genome assembled into two contigs, a 2.35 Mbp contig (M1404_1) and a 66 kb contig (M1404_2), despite having 360x depth across the genome. Most publicly available complete *P. multocida* genomes have under 200x depth, indicating that 360x depth should be enough depth to generate a closed genome.

Given strain M1404 has previously been shown to contain several prophages [2], we searched the genome for prophage regions using PHASTER, identifying six different prophage regions (Table S8). Five of the prophage regions were identified on the 2.35 Mbp contig, and one was identified on the 66 kb contig. The prophage region on the 66 kb contig and three prophage regions on the 2.35 Mbp contig shared >99% nucleotide identity, and were homologs of bacteriophage Mu, suggesting that the same prophage has inserted three times into the genome. The region flanking the prophage on the 66 kb contig matched genomic DNA from several other *P. multocida* strains. Mauve alignments showed the region flanking the prophage on the 66 kb contig was present as an intact segment in several other complete *P. multocida* genomes (Figure S9). Alignment of the 2.35 Mbp contig with the same genomes showed a gap where the expected genomic DNA sequence should be in the M1404 chromosome at the assembly gap (Fig S9). Furthermore, mapping and individual read-level analysis of the nanopore and Illumina reads to the 2.35 Mbp contig and the 66 kb contig showed that no single reads went further than assembly junction for either contig. Together these data suggest that the 66 kb contig is likely an excised prophage that has captured genomic DNA and may exist as a separate replicon in M1404. The 66 kb region contained six genes identified as essential for growth in rich media, which suggests this region cannot be lost. As we could not generate a closed genome, the M1404 assembly in two contigs was used for TraDIS analysis, with the 2.35 Mbp contig made to start at *dnaA*.

Supplemental Tables

**S1 Table**. Infectious dose and number of mice used for *P. multocida* systemic infections performed in this study

| Strain | Strain number | Mice used for experiment | Infectious dose in CFU^1^ |
| --- | --- | --- | --- |
| VP161 *Himar1* mutant library | NA^2^ | Two males and two females | 1.44 x 10^7^ |
| M1404 *Himar1* mutant library | NA | Two males and two females | 1.95 x 10^7^ |
| Wild-type M1404 harbouring empty vector | AL4221 | Three females | 3.30 x 10^4^ |
| Wild-type M1404 harbouring empty vector | AL4221 | Three males | 2.50 x 10^4^ |
| M1404 *alsT_1* mutant harbouring empty vector | AL4853 | Three females | 1.75 x 10^5^ |
| M1404 *alsT_1* mutant harbouring empty vector | AL4853 | Three males | 1.15 x 10^5^ |
| M1404 *crp* mutant harbouring empty vector | AL4855 | Three females | 1.70 x 10^5^ |
| M1404 *crp* mutant harbouring empty vector | AL4855 | Three males | 1.48 x 10^5^ |
| M1404 *cyaA* mutant harbouring empty vector | AL4857 | Three females | 1.87 x 10^5^ |
| M1404 *cyaA* mutant harbouring empty vector | AL4857 | Three males | 1.40 x 10^5^ |

^1^ CFU – Colony forming units

^2^ NA – Not applicable

**S2 Table.** TraDIS mapping statistics for the input transposon mutant libraries

| Library | Transposon-DNA junction reads | Mapped reads | Number of unique *Himar1* insertion sites (UIS^1^) | Resolution (Distance between UIS) |
| --- | --- | --- | --- | --- |
| Input libraries |  |  |  |  |
| VP161 |  |  |  |  |
| Library 1 | 1,757,473 | 1,211,338 | 42,771 | 52.51 |
| Library 2 | 1,764,365 | 1,240,274 | 42,803 | 52.47 |
| Library 3 | 1,757,577 | 1,222,848 | 42,518 | 52.83 |
| Library 4 | 1,764,661 | 1,218,068 | 42,060 | 53.40 |
| Total |  |  | 71,190 | 31.55 |
| M1404 |  |  |  |  |
| Library 1 | 6,981,977 | 5,807,837 | 38,852 | 62.30 |
| Library 2 | 6,648,491 | 5,590,017 | 37,424 | 64.68 |
| Library 3 | 7,569,978 | 6,337,831 | 39,834 | 60.77 |
| Library 4 | 6,784,647 | 5,698,351 | 38,252 | 63.28 |
| Total |  |  | 60,416 | 40.07 |
| Mice infected with VP161 |  |  |  |  |
| Blood |  |  |  |  |
| Mouse 1 | 1,770,340 | 1,313,856 | 40,159 | 55.93 |
| Mouse 2 | 1,734,955 | 1,254,692 | 40,924 | 54.88 |
| Mouse 3 | 1,775,359 | 1,298,576 | 38,852 | 57.81 |
| Mouse 4 | 1,767,708 | 1,302,992 | 39,768 | 56.48 |
| Total |  |  | 69,813 | 32.17 |
| Liver |  |  |  |  |
| Mouse 1 | 1,770,960 | 1,317,312 | 39,695 | 56.58 |
| Mouse 2 | 1,782,436 | 1,326,244 | 40,916 | 54.89 |
| Mouse 3 | 1,776,693 | 1,330,889 | 39,400 | 57.01 |
| Mouse 4 | 1,773,297 | 1,300,166 | 40,860 | 54.97 |
| Total |  |  | 71,057 | 31.61 |
| Spleen |  |  |  |  |
| Mouse 1 | 1,732,783 | 1,327,824 | 39,165 | 57.35 |
| Mouse 2 | 1,775,808 | 1,317,514 | 41,124 | 54.62 |
| Mouse 3 | 1,777,345 | 1,299,770 | 40,803 | 55.05 |
| Mouse 4 | 1,779,576 | 1,307,655 | 41,408 | 54.24 |
| Total |  |  | 73,400 | 30.60 |
| Mice infected with M1404 |  |  |  |  |
| Blood |  |  |  |  |
| Mouse 1 | 6,511,884 | 5,524,455 | 31,048 | 77.96 |
| Mouse 2 | 6,106,712 | 5,195,119 | 32,709 | 74.01 |
| Mouse 3 | 7,248,536 | 6,114,855 | 33,175 | 72.97 |
| Mouse 4 | 6,832,751 | 5,786,335 | 32,246 | 75.07 |
| Total |  |  | 56,761 | 42.65 |
| Liver |  |  |  |  |
| Mouse 1 | 7,954,019 | 6,748,110 | 31,770 | 76.19 |
| Mouse 2 | 5,330,169 | 4,430,316 | 30,228 | 80.08 |
| Mouse 3 | 9,066,531 | 7,847,303 | 36,144 | 66.97 |
| Mouse 4 | 6,347,637 | 5,32,2350 | 32,737 | 73.94 |
| Total |  |  | 56,948 | 42.51 |
| Spleen |  |  |  |  |
| Mouse 1 | 7,059,436 | 6,025,578 | 30,034 | 80.60 |
| Mouse 2 | 6,674,658 | 5,671,969 | 32,796 | 73.81 |
| Mouse 3 | 7,936,658 | 6,713,440 | 34,763 | 69.63 |
| Mouse 4 | 7,998,355 | 6,788,619 | 33,347 | 72.59 |
| Total |  |  | 57,371 | 42.19 |

^1^UIS – unique *Himar1* insertion sites

**S3 Table.** Genes in the 100% *P. multocida* core genome (1,564 genes) that encode proteins with matches to the curated database of *P. multocida* virulence factor and antibiotic resistance genes [1] at >75% amino acid identity.

| General function | Genes |
| --- | --- |
| Iron receptors | *hbpA*^1^, *hgbA*, PM0741, PM1081, PM1428 |
| Iron transporters | *tonB-exbBD*, *afuABC*, *afuA_2*, *afuA_3*, *fecCDE*, *fbpB*, *fbpC*, *yfeABCD* |
| Outer membrane proteins and fimbriae | *comE1*, *oma87*^1^, *ompH_2*, *ptfA* |
| Sialic acid uptake and utilisation | *nanP*, *nanU*, *nanB* |
| Methionine uptake | *metQ* |
| Transcriptional regulators, two component systems, and the stringent response | *spoT*^1^, *fis*, *fur*^1^, *hfq*, *qseB*, *qseC*, PM0442^1^ |
| Quorum sensing | *lsrABCDFGKR-luxS* |
| LPS biosynthesis | *hptA*, *hptB*, *hptC*, *hptD*, *lpt-3*, *gtcB*, *kdtA*^1^ |

^1^Identified as essential in VP161 and M1404 for growth in rich media

**S4 Table.** Genes that result in decreased *in vivo* fitness when disrupted in *P. multocida* strain VP161 or M1404, as identified by TraDIS analysis of mutants recovered from either the blood, liver or spleen following systemic infections in BALB/c mice.

|  |  |  |  |  | VP161 | | | | M1404 | | | |
| --- | --- | --- | --- | --- | --- | --- | --- | --- | --- | --- | --- | --- |
| VP161 locus tag^1^ | VP161 gene | M1404 locus tag^2^ | M1404 gene | Function | Rich media | Blood | Liver | Spleen | Rich media | Blood | Liver | Spleen |
| 0005 | *crp* | 00005 | *crp* | cAMP-activated global transcriptional regulator | Yes | NA^3^ | NA | NA | No | Yes | Yes | Yes |
| NA | *NA* | 00018 | *lex1* | Lipooligosaccharide biosynthesis protein | NA | NA | NA | NA | No | Yes | Yes | Yes |
| 0075 | *dus* | 00072 | *dusB* | tRNA-dihydrouridine synthase B | No | Yes | Yes | Yes | No | No | No | No |
| 0076 | *fis* | 00073 | *fis* | DNA-binding protein | No | Yes | Yes | Yes | No | NA | NA | NA |
| 0077 | *purL* | 00074 | *purL* | Phosphoribosylformylglycinamidine synthase | No | Yes | Yes | Yes | No | Yes | Yes | Yes |
| 0089 | *pgm* | 00086 | *pgm* | Phosphoglucomutase, capsule monomer biosynthesis | No | Yes | Yes | Yes | No | Yes | Yes | Yes |
| 0094 | 0094 | 00091 | 00091 | 47 kDa outer membrane protein | No | Yes | Yes | Yes | No | No | No | No |
| 0183 | 0183 | 00293 | 00293 | hypothetical protein | No | Yes | Yes | Yes | No | No | No | No |
| 0184 | *rnt* | 00294 | *rnt* | Ribonuclease T | Yes | No | No | No | No | Yes | No | No |
| 0191 | *nudB* | 00301 | *nudB* | Dihydroneopterin triphosphate diphosphatase | No | No | No | No | No | No | No | Yes |
| 0207 | *pal* | 00317 | *pal* | Peptidoglycan-associated lipoprotein | No | No | Yes | No | Yes | NA | NA | NA |
| 0219 | *holD* | 00327 | *holD* | DNA polymerase III subunit psi | No | No | No | No | No | Yes | No | No |
| 0242 | *purA* | 00350 | *purA* | Adenylosuccinate synthetase | No | Yes | Yes | Yes | No | Yes | Yes | Yes |
| 0252 | 0252 | 00360 | 00360 | tRNA-Met | Yes | NA | NA | NA | No | No | Yes | No |
| 0261 | *rpoZ* | 00369 | *rpoZ* | DNA-directed RNA polymerase subunit omega | No | No | No | No | No | No | No | Yes |
| 0281 | *hutZ_1* | 01072 | *hutZ_1* | Heme oxygenase | No | Yes | Yes | Yes | No | No | No | No |
| 0490 | 0490 | 00853 | 00853 | hypothetical protein | No | No | No | No | No | Yes | No | Yes |
| 0496 | 0496 | 00849 | 00849 | hypothetical protein | No | No | No | No | No | Yes | Yes | Yes |
| 0514 | *kdsC* | 00831 | *kdsC* | 3-deoxy-D-manno-octulosonate 8-phosphate phosphatase | Yes | NA | NA | NA | No | Yes | Yes | No |
| 0519 | *alsT_1* | 00826 | *alsT_1* | Amino-acid carrier protein | No | Yes | Yes | Yes | No | Yes | Yes | Yes |
| 0543 | *mdh* | 00805 | *mdh* | Malate dehydrogenase | No | No | No | No | No | Yes | No | No |
| 0549 | *prmC* | 00799 | *prmC* | Release factor glutamine methyltransferase | No | Yes | Yes | Yes | No | Yes | Yes | Yes |
| 0607 | *purE* | 00736 | *purE* | N5-carboxyaminoimidazole ribonucleotide mutase | No | Yes | Yes | Yes | No | Yes | Yes | Yes |
| 0608 | *purK* | 00735 | *purK* | N5-carboxyaminoimidazole ribonucleotide synthase | No | Yes | Yes | No | No | Yes | Yes | Yes |
| 0609 | *aspC* | 00734 | *aspC* | Aspartate aminotransferase | No | Yes | No | Yes | No | Yes | Yes | Yes |
| 0693 | *ubiX* | 00652 | *ubiX* | Flavin prenyltransferase | No | Yes | Yes | Yes | No | NA | NA | NA |
| 0694 | *purF* | 00651 | *purF* | Amidophosphoribosyltransferase | No | Yes | Yes | Yes | No | Yes | Yes | Yes |
| 0695 | *cvpA* | 00650 | *cvpA* | Colicin V production protein | No | Yes | Yes | Yes | No | No | No | Yes |
| 0727 | *tyrP_1* | 00619 | *tyrP_2* | Tyrosine-specific transport protein | No | Yes | Yes | Yes | No | No | No | No |
| 0773 | *phyB* | 00576 | *lipB* | Homology to capsule phospholipid substitution proteins | No | Yes | Yes | Yes | No | Yes | Yes | Yes |
| 0774 | *phyA* | NA | *NA* | Homology to capsule phospholipid substitution proteins | No | Yes | Yes | Yes | NA | NA | NA | NA |
| 0775 | *hyaE* | NA | *NA* | Hyaluronic acid capsule biosynthesis protein | No | Yes | Yes | Yes | NA | NA | NA | NA |
| 0776 | *hyaD* | NA | *NA* | Hyaluronic acid synthase | No | Yes | Yes | Yes | NA | NA | NA | NA |
| 0777 | *hyaC* | NA | *NA* | UDP-glucose 6-dehydrogenase | No | Yes | Yes | Yes | NA | NA | NA | NA |
| 0778 | *hyaB* | NA | *NA* | Hyaluronic acid capsule biosynthesis | No | Yes | Yes | Yes | NA | NA | NA | NA |
| 0779 | *hexD* | 00565 | *cexD* | Capsule export | Yes | Yes | Yes | No | No | Yes | Yes | Yes |
| 0780 | *hexC* | 00564 | *cexC* | Capsule export | Yes | Yes | Yes | Yes | No | Yes | Yes | Yes |
| 0781 | *hexB* | 00563 | *cexB* | Capsule transport protein | Yes | NA | NA | NA | No | Yes | Yes | Yes |
| 0782 | *hexA* | 00562 | *cexA* | Capsule transport ATP-binding protein | No | NA | NA | NA | No | Yes | Yes | Yes |
| NA | *NA* | 00566 | *lipA* | Capsule attachment | NA | NA | NA | NA | No | Yes | Yes | Yes |
| NA | *NA* | 00567 | *bcbI* | Capsule biosynthesis | NA | NA | NA | NA | No | Yes | Yes | Yes |
| NA | *NA* | 00568 | *bcbH* | Capsule biosynthesis | NA | NA | NA | NA | No | Yes | Yes | Yes |
| NA | *NA* | 00569 | *bcbG* | Capsule biosynthesis | NA | NA | NA | NA | No | Yes | Yes | Yes |
| NA | *NA* | 00570 | *bcbF* | Capsule biosynthesis | NA | NA | NA | NA | No | Yes | Yes | Yes |
| NA | *NA* | 00571 | *bcbE* | Capsule biosynthesis | NA | NA | NA | NA | No | Yes | Yes | Yes |
| NA | *NA* | 00572 | *bcbD* | Capsule biosynthesis | NA | NA | NA | NA | No | Yes | Yes | Yes |
| NA | *NA* | 00573 | *bcbC* | Capsule synthase | NA | NA | NA | NA | No | Yes | Yes | Yes |
| NA | *NA* | 00574 | *bcbB* | UDP-N-acetyl-D-mannosamine dehydrogenase | NA | NA | NA | NA | No | Yes | Yes | Yes |
| NA | *NA* | 00575 | *bcbA* | UDP-N-acetylglucosamine 2-epimerase | NA | NA | NA | NA | No | Yes | Yes | Yes |
| 0819 | *purC* | 00529 | *purC* | Phosphoribosylaminoimidazole-succinocarboxamide synthase | No | Yes | Yes | Yes | No | Yes | Yes | Yes |
| 0845 | *aroA* | 00506 | *aroA* | 3-phosphoshikimate 1-carboxyvinyltransferase | No | Yes | Yes | Yes | No | Yes | Yes | Yes |
| 0846 | *ubiG* | 00505 | *ubiG* | Ubiquinone biosynthesis O-methyltransferase | No | Yes | Yes | No | Yes | NA | NA | NA |
| 0872 | *folK* | 00431 | *folK* | 2-amino-4-hydroxy-6- hydroxymethyldihydropteridine pyrophosphokinase | Yes | NA | NA | NA | Yes | Yes | Yes | Yes |
| 0878 | 0878 | 00425 | 00425 | Putative Na/H antiporter | Yes | Yes | Yes | No | Yes | Yes | No | Yes |
| 0891 | *hldE* | 00412 | *hldE* | Bifunctional protein, LPS biosynthesis | No | Yes | Yes | Yes | No | Yes | Yes | Yes |
| 0914 | *hfq* | 00389 | *hfq* | RNA-binding protein | No | Yes | Yes | Yes | No | No | No | No |
| 0919 | *sapA* | 00384 | *sapA* | Peptide transport periplasmic protein | No | No | No | No | No | Yes | Yes | Yes |
| 0920 | *sapB_1* | 00383 | *sapB_1* | Putrescine export system permease protein | No | Yes | No | No | Yes | NA | NA | NA |
| 0921 | *sapC* | 00382 | *sapC* | Peptide transport system permease protein | Yes | No | Yes | No | Yes | NA | NA | NA |
| 0932 | *guaB* | 01083 | *guaB* | Inosine-5'-monophosphate dehydrogenase | Yes | NA | NA | NA | No | Yes | Yes | Yes |
| 0942 | *galE* | 01093 | *galE* | UDP-glucose 4-epimerase | No | No | No | Yes | No | Yes | Yes | Yes |
| 0994 | *trhP* | 01144 | *trhP* | tRNA wobble base hydroxylation protein | No | No | No | No | No | Yes | Yes | No |
| 1003 | *purD* | 01153 | *purD* | Phosphoribosylamine--glycine ligase | No | Yes | Yes | Yes | No | Yes | Yes | Yes |
| 1006 | *purH* | 01156 | *purH* | Bifunctional purine biosynthesis protein | No | Yes | Yes | Yes | No | Yes | Yes | Yes |
| 1029 | *epmA* | 01179 | *epmA* | Elongation factor P--(R)-beta-lysine ligase | No | Yes | Yes | Yes | No | Yes | No | No |
| 1044 | *neuA_1* | 01194 | *neuA_1* | N-acylneuraminate cytidylyltransferase | No | No | No | No | No | Yes | Yes | Yes |
| 1053 | *mlaB* | 01203 | *mlaB* | Phospholipid ABC transporter protein | Yes | No | No | No | No | Yes | No | No |
| 1055 | *mlaD* | 01205 | *mlaD* | Phospholipid ABC transporter-binding protein | No | No | No | No | No | Yes | No | No |
| 1056 | *mlaE* | 01206 | *mlaE* | Phospholipid ABC transporter permease protein | No | No | No | No | No | Yes | No | No |
| 1057 | *mlaF* | 01207 | *mlaF* | Phospholipid import ATP-binding protein | No | No | No | No | No | Yes | Yes | No |
| 1061 | *ptsN* | 01211 | *ptsN* | Nitrogen regulatory protein | No | No | No | No | Yes | No | Yes | No |
| 1068 | *ychF* | 01218 | *ychF* | Ribosome-binding ATPase | No | No | No | No | No | Yes | No | Yes |
| 1109 | *gmhA* | 01259 | *gmhA* | Phosphoheptose isomerase | No | Yes | Yes | Yes | No | Yes | Yes | Yes |
| 1136 | *epmB* | 01285 | *epmB* | L-lysine 2,3-aminomutase | No | Yes | Yes | Yes | No | No | No | No |
| 1172 | *lepA* | 01321 | *lepA* | Elongation factor 4 | Yes | No | No | No | Yes | No | No | Yes |
| 1217 | *purM* | 01364 | *purM* | Phosphoribosylformylglycinamidine cyclo-ligase | No | Yes | Yes | Yes | No | Yes | Yes | Yes |
| 1218 | *purN* | 01365 | *purN* | Phosphoribosylglycinamide formyltransferase | No | Yes | Yes | Yes | No | Yes | Yes | Yes |
| 1273 | *lon* | 01420 | *lon* | Lon protease | Yes | No | No | No | No | Yes | Yes | Yes |
| 1281 | *srlB* | 01428 | *srlB* | PTS system glucitol/sorbitol-specific EIIA component | No | No | No | No | No | Yes | Yes | Yes |
| 1294 | *mreB* | 01441 | *mreB* | Rod shape-determining protein | Yes | No | No | No | No | No | Yes | No |
| 1310 | *mrdA* | 01462 | *mrdA* | Peptidoglycan D,D-transpeptidase | No | No | No | No | Yes | Yes | Yes | No |
| 1321 | *plsX* | 01473 | *plsX* | Phosphate acyltransferase | No | Yes | Yes | Yes | Yes | NA | NA | NA |
| 1346 | *cpxA* | 01498 | *cpxA* | Sensor histidine kinase | No | No | No | No | No | Yes | No | Yes |
| 1359 | *rph* | 01511 | *rph* | Ribonuclease PH | No | No | No | No | No | Yes | No | Yes |
| 1376 | 1376 | 01528 | 01528 | putative ferredoxin-like protein | Yes | NA | NA | NA | No | Yes | No | No |
| 1383 | *purB* | 01535 | *purB* | Adenylosuccinate lyase | No | No | No | No | No | Yes | Yes | Yes |
| 1384 | *hflD* | 01536 | *hflD* | High frequency lysogenization protein | No | No | No | No | No | Yes | Yes | Yes |
| 1390 | *hptC* | 01542 | *hptC* | ADP-heptose--LPS heptosyltransferase-adds Hep II to Hep I in LPS | No | No | No | No | No | Yes | Yes | Yes |
| 1422 | *cyaA* | 01575 | *cyaA* | Adenylate cyclase | No | No | No | No | No | Yes | Yes | No |
| 1431 | *1431* | 01584 | *01584* | hypothetical protein | No | No | No | No | No | Yes | Yes | Yes |
| 1432 | *dsbA* | 01585 | *dsbA* | Thiol:disulfide interchange protein DsbA | No | No | No | No | No | No | No | Yes |
| 1480 | *tufA_1* | 01630 | *tufB_1* | Elongation factor Tu | No | No | No | No | No | Yes | Yes | Yes |
| 1572 | *gpsA* | 01723 | *gpsA* | Glycerol-3-phosphate dehydrogenase (NAD(P)+) | No | Yes | Yes | Yes | Yes | NA | NA | NA |
| 1604 | *pabA* | 01755 | *pabA* | Aminodeoxychorismate/anthranilate synthase component 2 | No | Yes | Yes | Yes | No | No | No | No |
| 1605 | *pabB* | 01756 | *pabB* | Aminodeoxychorismate synthase component 1 | No | Yes | Yes | Yes | No | No | No | No |
| NA | *NA* | 01811 | 01811 | Surface lipoprotein PmSLP-3 | NA | NA | NA | NA | No | Yes | Yes | Yes |
| 1663 | 1663 | 01812 | 01812 | Slam exporter | No | No | No | No | No | Yes | Yes | Yes |
| 1664 | *rsmD* | 01813 | *rsmD* | Ribosomal RNA small subunit methyltransferase D | No | No | No | No | No | Yes | No | Yes |
| 1720 | *asnC* | 01872 | *asnC* | Regulatory protein AsnC | No | Yes | Yes | Yes | No | Yes | Yes | Yes |
| 1721 | *asnA* | 01873 | *asnA* | Aspartate--ammonia ligase | No | Yes | Yes | Yes | No | Yes | Yes | Yes |
| 1798 | *thiE* | 01958 | *thiE* | Thiamine-phosphate synthase | Yes | NA | NA | NA | Yes | Yes | Yes | Yes |
| 1809 | *aroE* | 01969 | *aroE* | Shikimate dehydrogenase (NADP(+)) | No | Yes | Yes | Yes | Yes | NA | NA | NA |
| 1828 | *galU* | 01988 | *galU* | UTP--glucose-1-phosphate uridylyltransferase | No | Yes | Yes | Yes | No | Yes | Yes | Yes |
| 1833 | *hptD* | 01993 | *hptD* | Addition of Hep III to the inner core | No | No | No | No | No | Yes | Yes | Yes |
| 1844 | *hptA* | 02001 | *hptA* | Heptosyltransferase-adds Hep I to glycoform A LPS | No | Yes | Yes | No | No | Yes | Yes | Yes |
| 1848 | *gctB* | 02005 | *gctB* | Galactosyltransferase-adds to Hep I in LPS | No | No | No | No | No | Yes | Yes | Yes |
| 1886 | *hldD* | 02046 | *hldD* | ADP-L-glycero-D-manno-heptose-6-epimerase | No | Yes | No | Yes | No | Yes | Yes | Yes |
| 1902 | *tufA_2* | 02062 | *tufB_2* | Elongation factor Tu | No | No | No | No | No | Yes | Yes | Yes |
| 1919 | *plpB* | 02078 | *plpB* | putative D-methionine-binding lipoprotein | No | Yes | Yes | Yes | No | Yes | Yes | Yes |
| 1920 | *metP* | 02079 | *metP* | Methionine import system permease protein | No | Yes | Yes | Yes | No | Yes | Yes | Yes |
| 1921 | *metN* | 02080 | *metN* | Methionine import ATP-binding protein | No | Yes | Yes | Yes | No | No | Yes | No |
| 1922 | *gmhB* | 02081 | *gmhB* | D-glycero-beta-D-manno-heptose-1,7-bisphosphate 7-phosphatase | No | No | Yes | No | Yes | NA | NA | NA |
| 1931 | *ubiH* | 02090 | *ubiH* | 2-octaprenyl-6-methoxyphenol hydroxylase | No | Yes | No | No | No | No | No | No |
| 1945 | *nanE* | 02104 | *nanE* | Putative N-acetylmannosamine-6-phosphate 2-epimerase | Yes | No | No | No | No | Yes | Yes | No |
| 1947 | *nanP* | 02106 | *nanP* | Sialic acid-binding periplasmic protein | No | No | No | No | No | Yes | Yes | Yes |
| 1948 | *nanU* | 02107 | *nanU* | Sialic acid TRAP transporter permease protein | No | No | No | No | No | Yes | Yes | Yes |
| 1971 | *ubiE_3* | 02130 | *ubiE* | Ubiquinone/menaquinone biosynthesis C-methyltransferase UbiE | No | Yes | No | Yes | No | NA | NA | NA |
| 1986 | *serA* | 02145 | *serA* | D-3-phosphoglycerate dehydrogenase | No | Yes | Yes | Yes | No | No | No | No |
| 2000 | *serB* | 02159 | *serB* | Phosphoserine phosphatase | No | Yes | Yes | Yes | No | No | No | No |
| 2079 | *tonB* | 02253 | *tonB* | TonB-ExbBD system | No | Yes | Yes | No | No | Yes | Yes | Yes |
| 2080 | *exbD* | 02254 | *exbD* | TonB-ExbBD system | No | Yes | Yes | No | No | Yes | Yes | Yes |
| 2081 | *exbB* | 02255 | *exbB* | TonB-ExbBD system | No | Yes | Yes | No | No | Yes | Yes | Yes |

^1^The number represents the VP161 locus tag without the PmVP161_ prefix

^2^The number represents the M1404 locus tag without the M1404_ prefix

^3^NA – Not applicable

**S5 Table.** Genes that result in increased *in vivo* fitness when disrupted in *P. multocida* strain VP161 or M1404, as identified by TraDIS analysis of mutants recovered from either the blood, liver or spleen following systemic infections in BALB/c mice.

|  |  |  |  |  | VP161 | | | | M1404 | | | |
| --- | --- | --- | --- | --- | --- | --- | --- | --- | --- | --- | --- | --- |
| VP161 locus tag^1^ | VP161 gene | M1404 locus tag^2^ | M1404 gene | Function | Rich media | Blood | Liver | Spleen | Rich media | Blood | Liver | Spleen |
| 0233 | 0233 | 00341 | 00341 | Hypothetical protein | No | No | No | No | No | Yes | No | No |
| 0256 | 0256 | 00364 | 00364 | Long-chain-fatty-acid--CoA ligase FadD15 | No | No | Yes | Yes | No | No | No | No |
| 0818 | *yeeZ* | 00530 | *yeeZ* | Protein YeeZ | Yes | Yes | Yes | Yes | Yes | Yes | Yes | Yes |
| NA^3^ | NA | 00860 | 00860 | Hypothetical protein | NA | NA | NA | NA | No | No | Yes | No |
| 0959 | *proQ* | 01109 | *proQ* | RNA chaperone | Yes | NA | NA | Yes | Yes | NA | NA | NA |
| 1146 | *yacG* | 01295 | *yacG* | DNA gyrase inhibitor | Yes | NA | NA | NA | Yes | Yes | No | No |
| 1492 | *gpt* | 01642 | *gpt* | Xanthine phosphoribosyltransferase | Yes | Yes | Yes | Yes | Yes | NA | NA | NA |
| 1514 | *speB* | 01672 | *speB* | Agmatinase | Yes | Yes | Yes | Yes | Yes | Yes | Yes | Yes |
| 1565 | 1565 | 01716 | 01716 | Hypothetical protein | No | No | No | No | No | Yes | Yes | Yes |
| 1804 | *ssuB* | 01964 | *ssuB* | Aliphatic sulfonates import ATP-binding protein | Yes | No | Yes | No | No | No | No | No |
| 1863 | *mltC* | 02022 | *mltC* | Membrane-bound lytic murein transglycosylase C | Yes | Yes | Yes | Yes | Yes | NA | NA | NA |
| 2098 | *rraA* | 02274 | *rraA* | Regulator of ribonuclease activity A | No | No | No | No | No | Yes | Yes | Yes |

^1^The number represents the VP161 locus tag without the PmVP161_ prefix

^2^The number represents the M1404 locus tag without the M1404_ prefix

^3^NA – Not applicable

**S6 Table.** Strains and plasmids used in this study.

| Strain or plasmid | Description | Source or reference |
| --- | --- | --- |
| Strains |  |  |
| *P. multocida* |  |  |
| M1404 | Bison haemorrhagic septicaemia isolate, serotype B:L2 | K. R. Rhoades, National Animal Disease Center, Ames, Iowa |
| VP161 | Chicken fowl cholera isolate, serotype A:1 | [3] |
| VP161-Tn*7* | VP161 with Tn*7* insertion downstream of *glmS*; Kan^R^ | [4] |
| AL2188 | VP161 harbouring pAL99S; Spec^R^ | [5] |
| AL4221 | M1404 harbouring pAL99S; Spec^R^ | This study |
| AL4853 | M1404 *alsT_1* insertional mutant harbouring pAL99S; Kan^R^ Spec^R^ | This study |
| AL4855 | M1404 *crp* insertional mutant harbouring pAL99S; Kan^R^ Spec^R^ | This study |
| AL4857 | M1404 *cyaA* insertional mutant harbouring pAL99S; Kan^R^ Spec^R^ | This study |
| AL4870 | M1404 *alsT_1* insertional mutant harbouring pAL2045; Kan^R^ Spec^R^ | This study |
| AL4871 | M1404 *cyaA* insertional mutant harbouring pAL2046; Kan^R^ Spec^R^ | This study |
| AL4872 | VP161 *cyaA* insertional mutant harbouring pAL99S; Kan^R^ Spec^R^ | This study |
| AL4873 | VP161 *cyaA* insertional mutant harbouring pAL2047; Kan^R^ Spec^R^ | This study |
| AL4874 | M1404 *crp* insertional mutant harbouring pAL2048; Kan^R^ Spec^R^ | This study |
| *E. coli* |  |  |
| DH5α | F^-^ *deoR endA1* *gyrA96 hsdR17* (r_K_^-^ m_K_^-^) *recA1 relA1* *supE44 thi-1* Δ(*lacZYAargFV169*) ᶲ80*lacZ*ΔM15 | Bethesda Research Laboratories |
| JKE201 | MFDpir Δ*mcrA* Δ(*mrr-hsdRMS-mcrBC*) *aac(3)IV*::*lacI^q^*; Erm^R^ | [6] |
| AL1296 | DH5α harbouring pAL99S; Spec^R^ | [5] |
| AL1995 | DH5α harbouring pAL953; Kan^R^ Spec^R^ | [5] |
| AL4487 | JKE201 harbouring pAL614; Amp^R^ Spec^R^ | This study |
| Plasmids |  |  |
| pAL99S | *P. multocida­*-*E. coli* expression plasmid; Spec^R^ P*tpiA* | [5] |
| pAL614 | RP4 mobilisable plasmid containing *Himar1*::Spec; RP4 Mob^+^ oriR6K Amp^R^ Spec^R^ tnpC9 |  |
| pAL953 | *P. multocida* vector containing ClosTron group II intron; Spec^R^ Kan^R^ | [5] |
| pAL2019 | pAL953 with the group II intron targeted to *alsT_1*; Kan^R^ Spec^R^ | This study |
| pAL2022 | pAL953 with the group II intron targeted to *crp*; Kan^R^ Spec^R^ | This study |
| pAL2023 | pAL953 with the group II intron targeted to *cyaA*; Kan^R^ Spec^R^ | This study |
| pAL2045 | Wild-type copy of *alsT_1* from M1404 cloned into pAL99S; Spec^R^ | This study |
| pAL2046 | Wild-type copy of *cyaA* from M1404 cloned into pAL99S; Spec^R^ | This study |
| pAL2047 | Wild-type copy of *cyaA* from VP161 cloned into pAL99S; Spec^R^ | This study |
| pAL2048 | Wild-type copy of *crp* from M1404 cloned into pAL99S; Spec^R^ | This study |

**S7 Table.** Oligonucleotides used in this study

| Oligonucleotide | Sequence (5’ – 3’)^1^ | Description |
| --- | --- | --- |
| Oligonucleotides used for TraDIS library production | | |
| BAP8034 | P‑G*ATCGGAAGAGCGGTTCAGCAGGTTTTTTTTTTCAAAAAAA*A | Splinkerette adapter top strand [7] |
| BAP8035 | G*AGATCGGTCTCGGCATTCCTGCTGAACCGCTCTTCCGATC*T | Splinkerette adapter bottom strand [7] |
| BAP8037 | CAAGCAGAAGACGGCATACGAGAT**TAAGGCGA**GAGATCGGTCTCGGCATTCC | Adapter-specific oligonucleotide used for TraDIS library amplification, contains an index sequence (in bold) and an Illumina P7 sequence at the 5’ |
| BAP8038 | CAAGCAGAAGACGGCATACGAGAT**CGTACTAG**GAGATCGGTCTCGGCATTCC | Adapter-specific oligonucleotide used for TraDIS library amplification, contains an index sequence (in bold) and an Illumina P7 sequence at the 5’ |
| BAP8039 | CAAGCAGAAGACGGCATACGAGAT**AGGCAGAA**GAGATCGGTCTCGGCATTCC | Adapter-specific oligonucleotide used for TraDIS library amplification, contains an index sequence (in bold) and an Illumina P7 sequence at the 5’ |
| BAP8040 | CAAGCAGAAGACGGCATACGAGAT**TCCTGAGC**GAGATCGGTCTCGGCATTCC | Adapter-specific oligonucleotide used for TraDIS library amplification, contains an index sequence (in bold) and an Illumina P7 sequence at the 5’ |
| BAP8042 | GGTTCTAGAGACCGGGGACTTATCAGC | Custom *Himar1* transposon-specific Illumina sequencing oligonucleotide |
| BAP8043 | TTCAGCAGGAATGCCGAGACCGATCTC | Custom adapter index-specific Illumina sequencing oligonucleotide |
| BAP8335 | CAAGCAGAAGACGGCATACGAGAT**GGACTCCT**GAGATCGGTCTCGGCATTCC | Adapter-specific oligonucleotide used for TraDIS library amplification, contains an index sequence (in bold) and an Illumina P7 sequence at the 5’ |
| BAP8336 | CAAGCAGAAGACGGCATACGAGAT**TAGGCATG**GAGATCGGTCTCGGCATTCC | Adapter-specific oligonucleotide used for TraDIS library amplification, contains an index sequence (in bold) and an Illumina P7 sequence at the 5’ |
| BAP8337 | CAAGCAGAAGACGGCATACGAGAT**CTCTCTAC**GAGATCGGTCTCGGCATTCC | Adapter-specific oligonucleotide used for TraDIS library amplification, contains an index sequence (in bold) and an Illumina P7 sequence at the 5’ |
| BAP8338 | CAAGCAGAAGACGGCATACGAGAT**CAGAGAGG**GAGATCGGTCTCGGCATTCC | Adapter-specific oligonucleotide used for TraDIS library amplification, contains an index sequence (in bold) and an Illumina P7 sequence at the 5’ |
| BAP8339 | CAAGCAGAAGACGGCATACGAGAT**GCTACGCT**GAGATCGGTCTCGGCATTCC | Adapter-specific oligonucleotide used for TraDIS library amplification, contains an index sequence (in bold) and an Illumina P7 sequence at the 5’ |
| BAP8350 | AATGATACGGCGACCACCGAGATCTACACCTCTAGAAAGTATAGGAACTTCGAACCG | *Himar1*-specific oligonucleotide for TraDIS library amplification, contains Illumina P5 at the 5’ end |
| BAP8356 | CAAGCAGAAGACGGCATACGAGAT**CGAGGCTG**GAGATCGGTCTCGGCATTCC | Adapter-specific oligonucleotide used for TraDIS library amplification, contains an index sequence (in bold) and an Illumina P7 sequence at the 5’ |
| BAP8357 | CAAGCAGAAGACGGCATACGAGAT**AAGAGGCA**GAGATCGGTCTCGGCATTCC | Adapter-specific oligonucleotide used for TraDIS library amplification, contains an index sequence (in bold) and an Illumina P7 sequence at the 5’ |
| BAP8358 | CAAGCAGAAGACGGCATACGAGAT**GTAGAGGA**GAGATCGGTCTCGGCATTCC | Adapter-specific oligonucleotide used for TraDIS library amplification, contains an index sequence (in bold) and an Illumina P7 sequence at the 5’ |
| BAP9946 | CAAGCAGAAGACGGCATACGAGAT**ATCACGAC**GAGATCGGTCTCGGCATTCC | Adapter-specific oligonucleotide used for TraDIS library amplification, contains an index sequence (in bold) and an Illumina P7 sequence at the 5’ |
| BAP9947 | CAAGCAGAAGACGGCATACGAGAT**ACAGTGGT**GAGATCGGTCTCGGCATTCC | Adapter-specific oligonucleotide used for TraDIS library amplification, contains an index sequence (in bold) and an Illumina P7 sequence at the 5’ |
| BAP9948 | CAAGCAGAAGACGGCATACGAGAT**ACCCAGCA**GAGATCGGTCTCGGCATTCC | Adapter-specific oligonucleotide used for TraDIS library amplification, contains an index sequence (in bold) and an Illumina P7 sequence at the 5’ |
| BAP10005 | CAAGCAGAAGACGGCATACGAGAT**CAGATCCA**GAGATCGGTCTCGGCATTCC | Adapter-specific oligonucleotide used for TraDIS library amplification, contains an index sequence (in bold) and an Illumina P7 sequence at the 5’ |
| BAP10006 | CAAGCAGAAGACGGCATACGAGAT**ACAAACGG**GAGATCGGTCTCGGCATTCC | Adapter-specific oligonucleotide used for TraDIS library amplification, contains an index sequence (in bold) and an Illumina P7 sequence at the 5’ |
| BAP10007 | CAAGCAGAAGACGGCATACGAGAT**AACCCCTC**GAGATCGGTCTCGGCATTCC | Adapter-specific oligonucleotide used for TraDIS library amplification, contains an index sequence (in bold) and an Illumina P7 sequence at the 5’ |
| BAP10008 | CAAGCAGAAGACGGCATACGAGAT**CCCAACCT**GAGATCGGTCTCGGCATTCC | Adapter-specific oligonucleotide used for TraDIS library amplification, contains an index sequence (in bold) and an Illumina P7 sequence at the 5’ |
| BAP10009 | CAAGCAGAAGACGGCATACGAGAT**CACCACAC**GAGATCGGTCTCGGCATTCC | Adapter-specific oligonucleotide used for TraDIS library amplification, contains an index sequence (in bold) and an Illumina P7 sequence at the 5’ |
| BAP10010 | CAAGCAGAAGACGGCATACGAGAT**GAAACCCA**GAGATCGGTCTCGGCATTCC | Adapter-specific oligonucleotide used for TraDIS library amplification, contains an index sequence (in bold) and an Illumina P7 sequence at the 5’ |
| BAP10011 | CAAGCAGAAGACGGCATACGAGAT**TGTGACCA**GAGATCGGTCTCGGCATTCC | Adapter-specific oligonucleotide used for TraDIS library amplification, contains an index sequence (in bold) and an Illumina P7 sequence at the 5’ |
| BAP10012 | CAAGCAGAAGACGGCATACGAGAT**AGGGTCAA**GAGATCGGTCTCGGCATTCC | Adapter-specific oligonucleotide used for TraDIS library amplification, contains an index sequence (in bold) and an Illumina P7 sequence at the 5’ |
| BAP10013 | CAAGCAGAAGACGGCATACGAGAT**AGGAGTGG**GAGATCGGTCTCGGCATTCC | Adapter-specific oligonucleotide used for TraDIS library amplification, contains an index sequence (in bold) and an Illumina P7 sequence at the 5’ |
| BAP10014 | CAAGCAGAAGACGGCATACGAGAT**TAGATCGC**GAGATCGGTCTCGGCATTCC | Adapter-specific oligonucleotide used for TraDIS library amplification, contains an index sequence (in bold) and an Illumina P7 sequence at the 5’ |
| BAP10015 | CAAGCAGAAGACGGCATACGAGAT**CTCTCTAT**GAGATCGGTCTCGGCATTCC | Adapter-specific oligonucleotide used for TraDIS library amplification, contains an index sequence (in bold) and an Illumina P7 sequence at the 5’ |
| BAP10016 | CAAGCAGAAGACGGCATACGAGAT**TATCCTCT**GAGATCGGTCTCGGCATTCC | Adapter-specific oligonucleotide used for TraDIS library amplification, contains an index sequence (in bold) and an Illumina P7 sequence at the 5’ |
| BAP10017 | CAAGCAGAAGACGGCATACGAGAT**AGAGTAGA**GAGATCGGTCTCGGCATTCC | Adapter-specific oligonucleotide used for TraDIS library amplification, contains an index sequence (in bold) and an Illumina P7 sequence at the 5’ |
| BAP10018 | CAAGCAGAAGACGGCATACGAGAT**GTAAGGAG**GAGATCGGTCTCGGCATTCC | Adapter-specific oligonucleotide used for TraDIS library amplification, contains an index sequence (in bold) and an Illumina P7 sequence at the 5’ |
| BAP10019 | CAAGCAGAAGACGGCATACGAGAT**ACTGCATA**GAGATCGGTCTCGGCATTCC | Adapter-specific oligonucleotide used for TraDIS library amplification, contains an index sequence (in bold) and an Illumina P7 sequence at the 5’ |
| BAP10020 | CAAGCAGAAGACGGCATACGAGAT**AAGGAGTA**GAGATCGGTCTCGGCATTCC | Adapter-specific oligonucleotide used for TraDIS library amplification, contains an index sequence (in bold) and an Illumina P7 sequence at the 5’ |
| BAP10021 | CAAGCAGAAGACGGCATACGAGAT**CTAAGCCT**GAGATCGGTCTCGGCATTCC | Adapter-specific oligonucleotide used for TraDIS library amplification, contains an index sequence (in bold) and an Illumina P7 sequence at the 5’ |
| ClosTron mutagenesis | | |
| BAP6544 | CGAAATTAGAAACTTGCGTTCAGTAAAC | EBS universal primer, used to retarget the group II intron to a specific target and for Sanger sequencing to confirm a single insertion |
| BAP10089 | AAAAAAGCTTATAATTATCCTTAGTGGCCGTCCAGGTGCGCCCAGATAGGGTG | IBS ClosTron oligonucleotide for retargeting the group II intron targeting region to *alsT_1* |
| BAP10090 | CAGATTGTACAAATGTGGTGATAACAGATAAGTCGTCCAGCCTAACTTACCTTTCTTTGT | EBS1d ClosTron oligonucleotide for retargeting the group II intron targeting region to *alsT_1* |
| BAP10091 | TGAACGCAAGTTTCTAATTTCGATTGCCACTCGATAGAGGAAAGTGTCT | EBS2 ClosTron oligonucleotide for retargeting the group II intron targeting region to *alsT_1* |
| BAP10109 | AAAAAAGCTTATAATTATCCTTAGGCAACGAAATGGTGCGCCCAGATAGGGTG | IBS ClosTron oligonucleotide for retargeting the group II intron targeting region to *crp* |
| BAP10110 | CAGATTGTACAAATGTGGTGATAACAGATAAGTCGAAATGATTAACTTACCTTTCTTTGT | EBS1d ClosTron oligonucleotide for retargeting the group II intron targeting region to *crp* |
| BAP10111 | TGAACGCAAGTTTCTAATTTCGATTTTGCCTCGATAGAGGAAAGTGTCT | EBS2 ClosTron oligonucleotide for retargeting the group II intron targeting region to *crp* |
| BAP10112 | AAAAAAGCTTATAATTATCCTTAAATCTCCCCGGCGTGCGCCCAGATAGGGTG | IBS ClosTron oligonucleotide for retargeting the group II intron targeting region to *cyaA* |
| BAP10113 | CAGATTGTACAAATGTGGTGATAACAGATAAGTCCCCGGCTATAACTTACCTTTCTTTGT | EBS1d ClosTron oligonucleotide for retargeting the group II intron targeting region to *cyaA* |
| BAP10114 | TGAACGCAAGTTTCTAATTTCGGTTAGATTCCGATAGAGGAAAGTGTCT | EBS2 ClosTron oligonucleotide for retargeting the group II intron targeting region to *cyaA* |
| Complementation plasmid production | | |
| BAP10092 | AAAAAACCCGGGAGGAGGAATAATGAGCATATTTTCTAC | Forward oligonucleotide flanking *alsT_1*, contains an *Xma*I restriction site |
| BAP10093 | AAAAAAAAGCTTTTAAGACCAAATATCGTTATCGAC | Reverse oligonucleotide flanking *alsT_1*, contains an *Hin*dIII restriction site |
| BAP10115 | AAAAAAGGATCCGTGAGTGGAATATCATTTGAATTAC | Forward oligonucleotide flanking *cyaA*, contains an *Bam*HI restriction site |
| BAP10116 | AAAAAAAAGCTTTTATGACATCGCTAATCGACTG | Reverse oligonucleotide flanking *cyaA*, contains an *Hin*dIII restriction site |
| BAP10117 | AAAAAAGGATCCATGGAGGTCTTCCGTGCAAG | Forward oligonucleotide flanking *crp*, contains an *Bam*HI restriction site |
| BAP10118 | AAAAAAAAGCTTGATTATCTTGTACCGTAAACGAC | Reverse oligonucleotide flanking *crp*, contains an *Hin*dIII restriction site |

^1^P- represents a 5’ phosphate and * represents a phosphorothioate bond

**S8 Table.** Prophage regions identified in the *de novo* assembled M1404 genome

| Region | Length | PHASTER score | Coding sequences | Genome position in nucleotides | Homolog | GC % |
| --- | --- | --- | --- | --- | --- | --- |
| M1404_1 |  |  |  |  |  |  |
| 1 | 73.5Kb | 150 | 99 | 95,238-168,755 | NC_000929 | 42.35 |
| 2 | 33.5Kb | 150 | 48 | 222,212-255,789 | NC_027382 | 42.44 |
| 3 | 36.7Kb | 150 | 49 | 403,519-440,233 | NC_027382 | 42.11 |
| 4 | 15.6Kb | 110 | 18 | 2,243,343-2,259,040 | NC_031940 | 38.06 |
| M1404_2 |  |  |  |  |  |  |
| 5 | 34.5Kb | 150 | 51 | 24,192-58,714 | NC_000929 | 42.36 |

Supplemental Figures

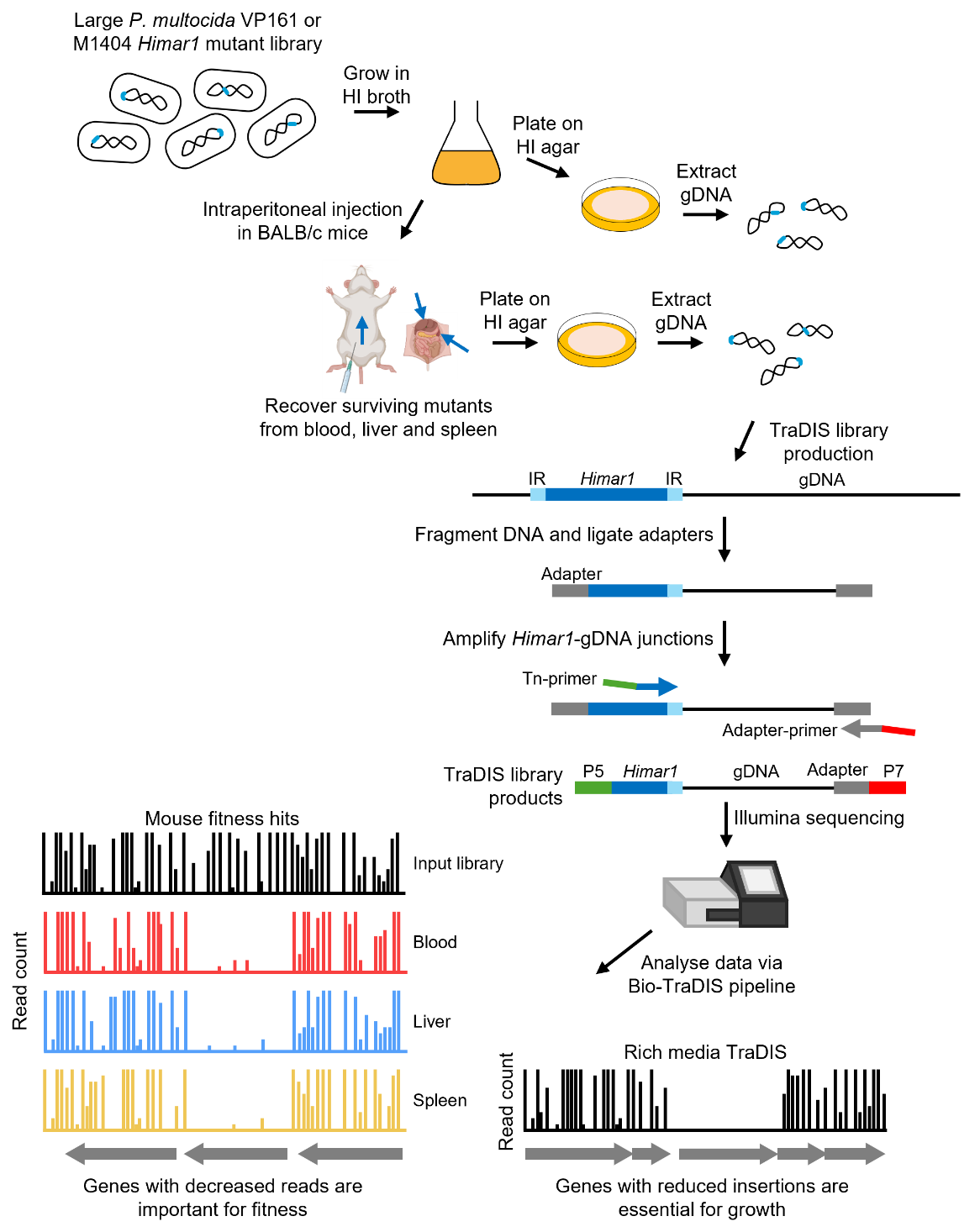

**S1 Fig.** Overview of the TraDIS methodology from this study. A large *P. multocida* strain M1404 *Himar1* mutant library was generated via conjugation. The M1404 *Himar1* library, along with a previously produced VP161 *Himar1* mutant library, were used to identify genes essential for growth in rich medium and also to perform systemic infections in BALB/c mice. Surviving mutants were recovered from the blood, liver, and spleen from mice, and plated onto heart infusion agar. Mutants were recovered, and genomic DNA (gDNA) extracted to produce TraDIS libraries. The gDNA was fragmented by sonication, end-repaired, and adapters ligated onto all fragments. Transposon-chromosome junctions were amplified using oligonucleotides specific to the *Himar1* inverted repeat and adapter, with oligonucleotides containing Illumina P5 and P7 sequences. The TraDIS libraries were then sequenced using an Illumina NextSeq, with data analysed using the Bio-TraDIS toolkit and related scripts. The normalized number of unique insertion sites per gene was compared to identify genes important for growth in rich media. Normalised read counts per gene were compared between the rich media TraDIS libraries, and each of the bloodstream, liver, and spleen TraDIS libraries to identify genes that result in either a fitness cost or benefit when disrupted by transposon insertions.

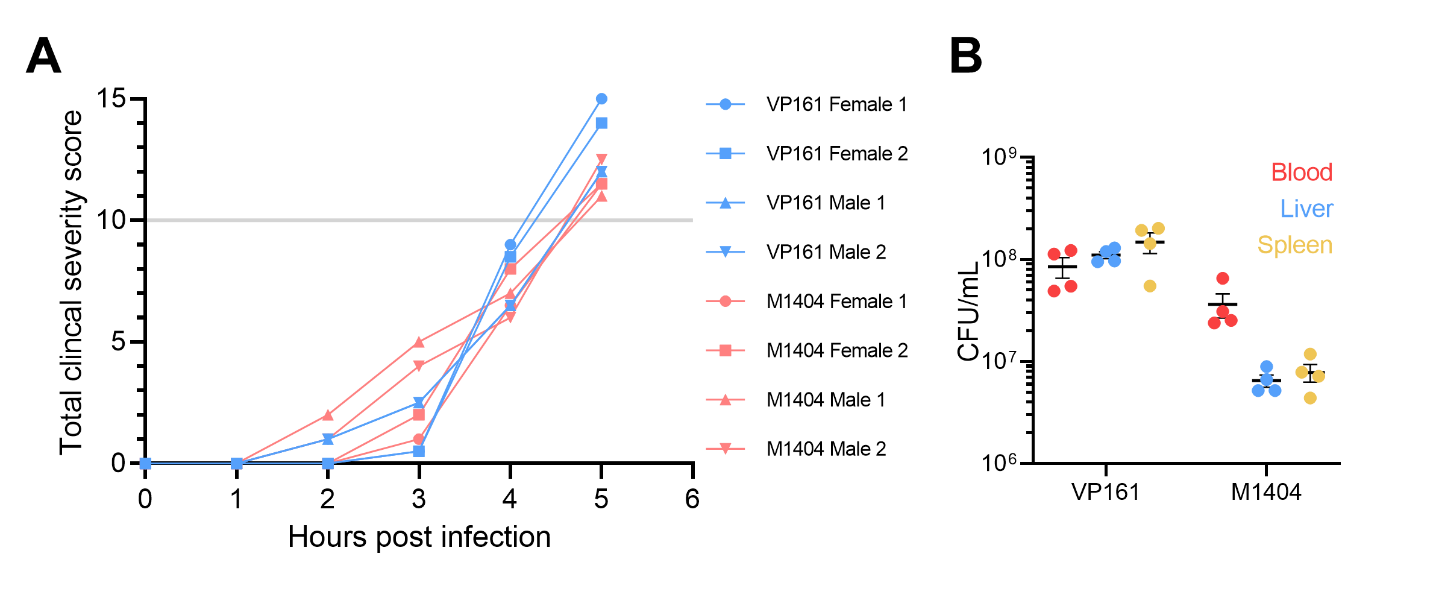

**S2 Fig.** Mouse systemic infections with the *P. multocida* strain VP161 and M1404 *Himar1* mutant libraries. For each library, two male and two female 6-10-week-old BALB/c mice were injected intraperitoneally with ~ 2 x 10^7^ CFU. **A.** Mice were monitored for clinical signs of systemic infection, with mice reaching the humane endpoint when they had a total score above 10 (grey line). **B.** Surviving VP161 or M1404 *Himar1* mutants were recovered from the bloodstream, liver, and spleen. Blood was resuspended in a total volume of 1 mL, and liver and spleen samples were homogenized in 1 mL of 1 x PBS, before being plated onto heart infusion agar.

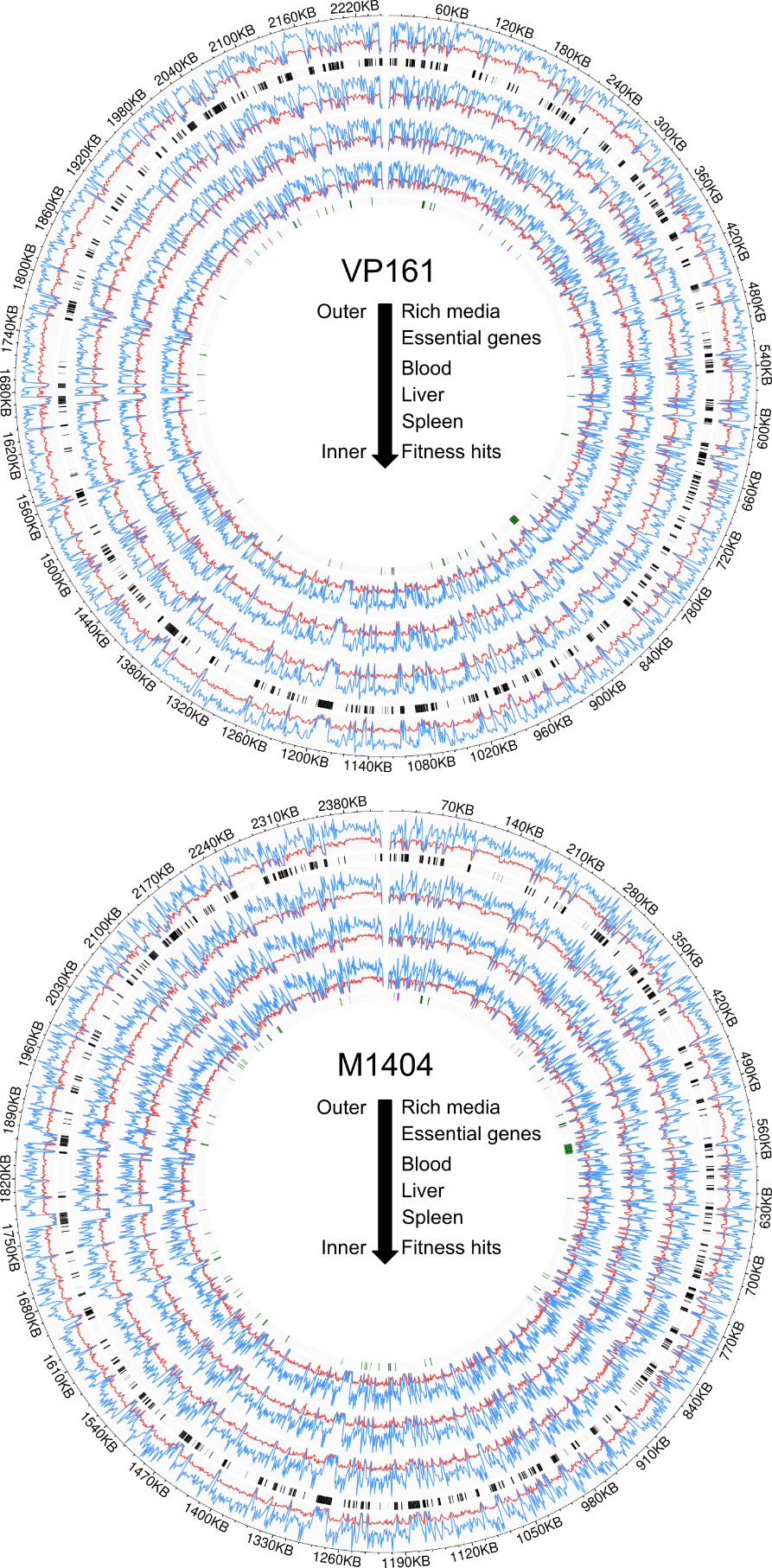

**S3 Fig.** Log_2_ read count (blue) and unique insertion sites (red) per 1 kb across the *P. multocida* VP161 and M1404 genome. Genes identified as essential for growth in rich media are shown as black bars in the second from outer ring, while fitness-cost genes are shown as green bars and fitness-benefit genes shown as purple bars in the inner ring.

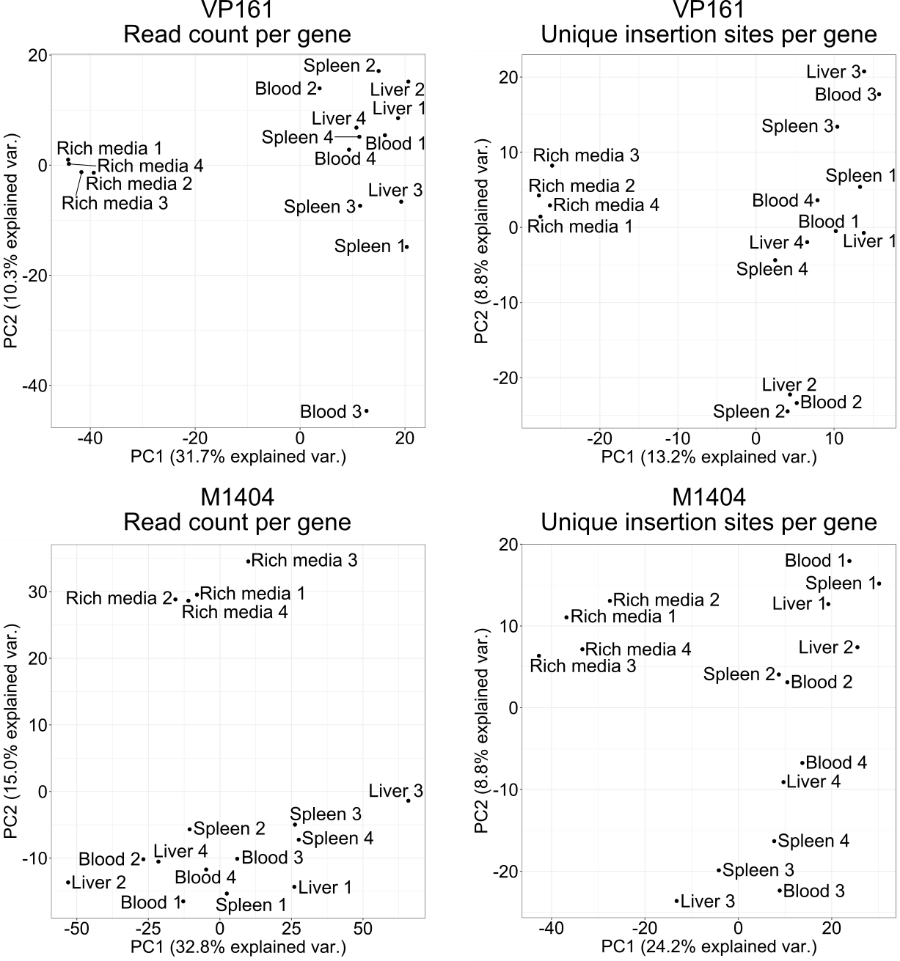

**S4 Fig.** Principal component analysis (PCA) plots for all VP161 and M1404 TraDIS libraries in this study. PCA was performed using prcomp in R, using either the number of unique insertion sites per gene or the number of reads per gene.

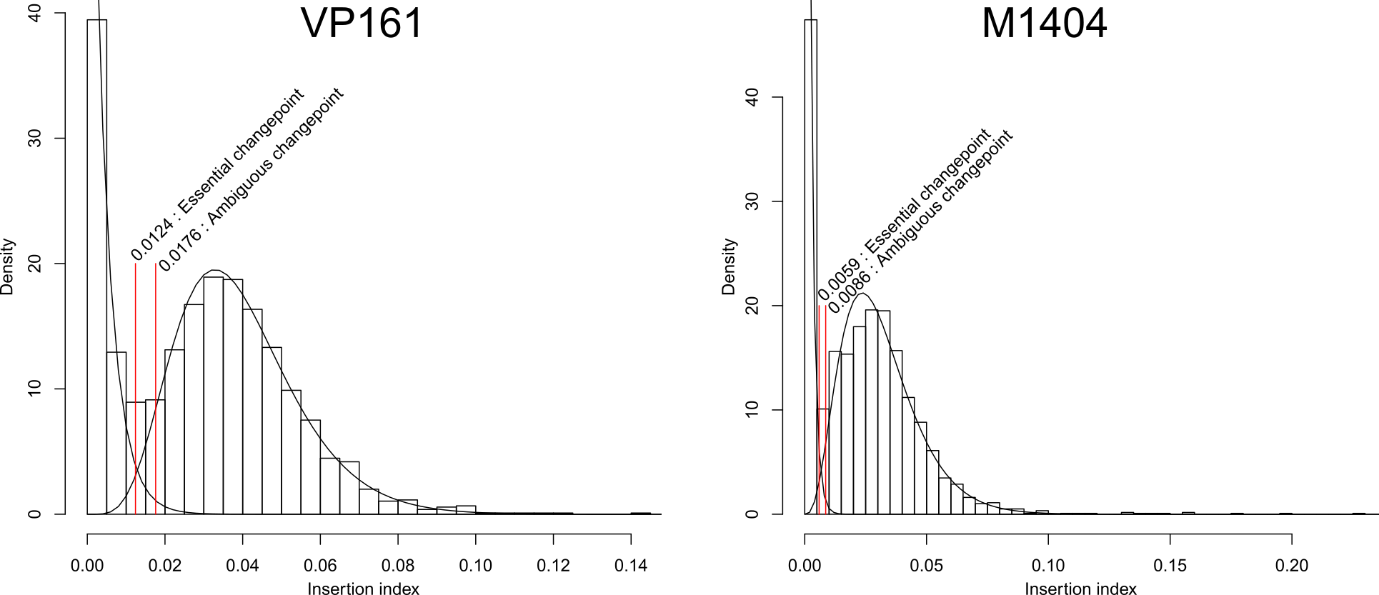

**S5 Fig.** Histograms of unique insertion sites (UIS) per gene and identification of essential gene cut-offs in the *P. multocida* strain VP161 and M1404 rich media TraDIS libraries. The number of UIS is divided by gene length to identify the insertion index, a normalized UIS count per gene. Histograms are generated using the insertion index from all genes, generating a bimodal distribution of essential and non-essential genes. Normal curves are drawn for the two sets, and the intersect between the normal curves is taken as the cut-off for gene essentiality.

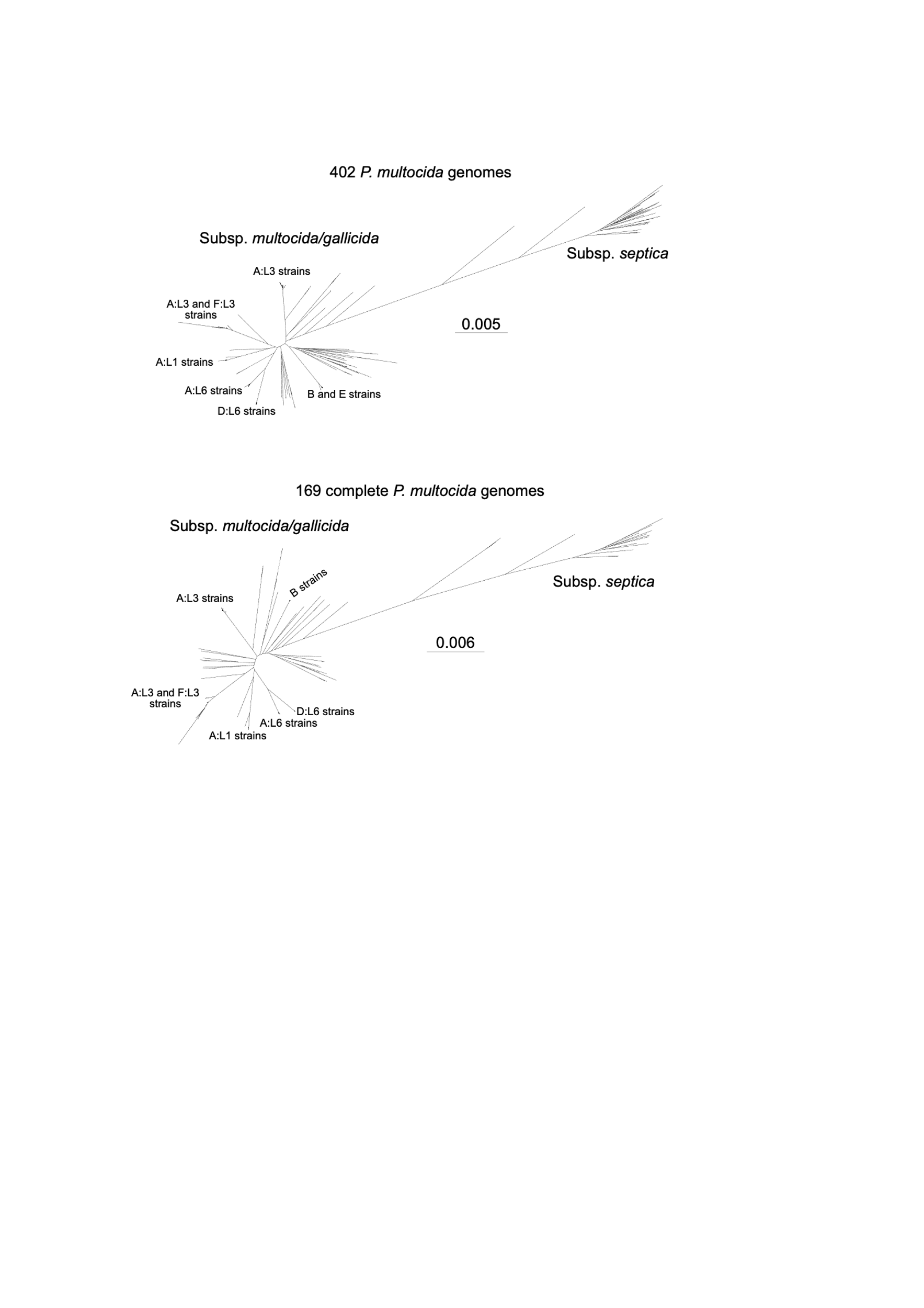

**S6 Fig.** Maximum-likelihood core-genome phylogeny of the 169 complete *P. multocida* genomes used to identify the 100% *P. multocida* core genome. Roary was used to generate a core-genome alignment using 850,282 sites, and IQ-TREE was used to generate the phylogeny (using model GTR+F+R10), with 1,000 bootstrap replicates (see supplemental material 6 for bootstrap values). Scale bar represents number of nucleotide substitutions per site. The maximum-likelihood core-genome phylogeny of 402 *P. multocida* strains, including incomplete genomes, was included as a comparison.

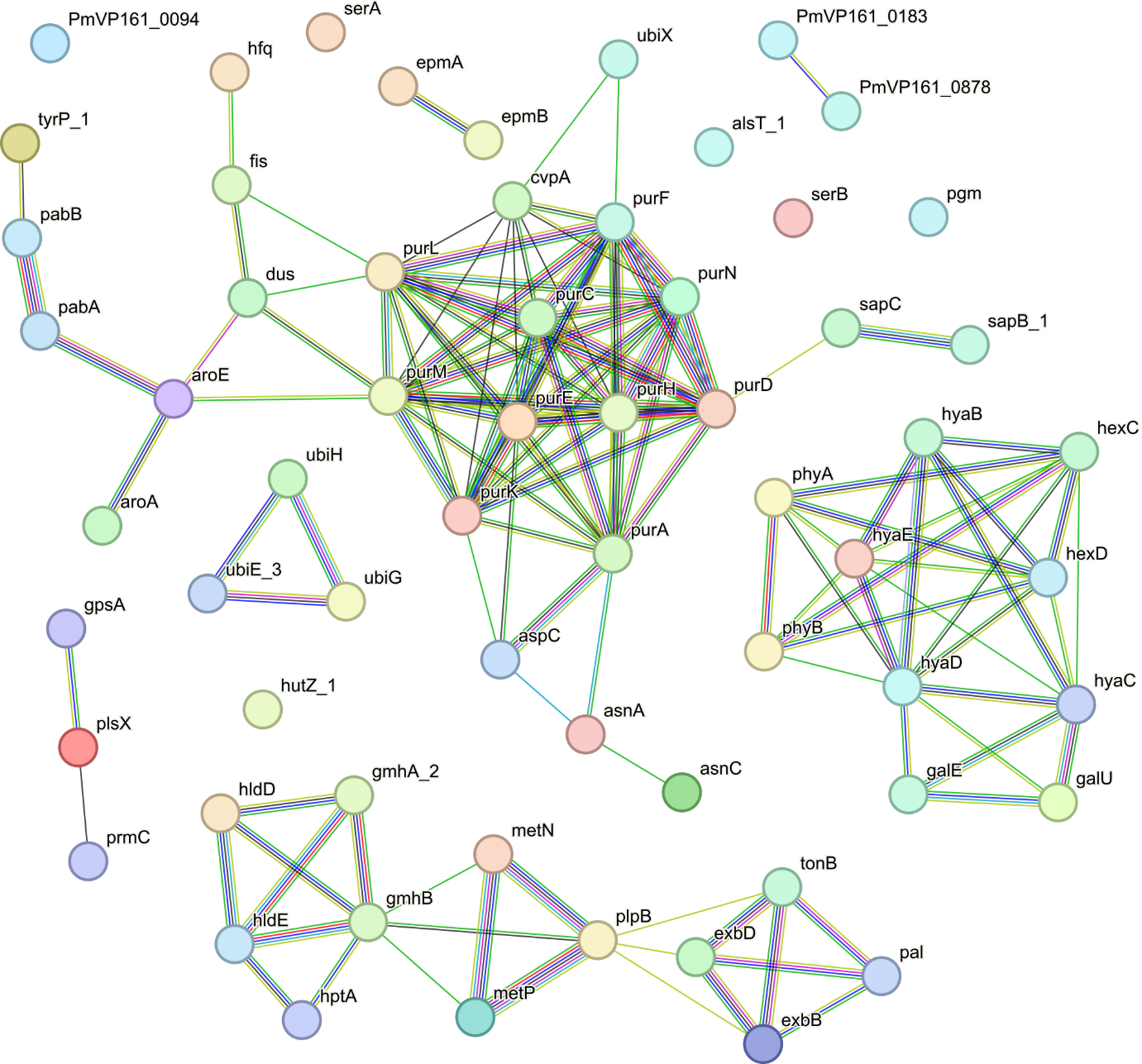

**S7 Fig.** STRING interaction map for *P. multocida* strain VP161 *in vivo* fitness genes. Genes were included if they were identified as important for VP161 survival in the bloodstream, liver, or spleen of BALB/c mice. The protein sequences of *in vivo* fitness genes were matched to *P. multocida* strain Pm70 in string, with all 63 genes matching to a Pm70 protein. The lines between genes show the type of evidence for interaction; red line - fusion evidence, green line - neighbourhood evidence, blue line - co-occurrence evidence, purple line - experimental evidence, yellow line - text-mining evidence, light blue line - database evidence, black line - co-expression evidence.

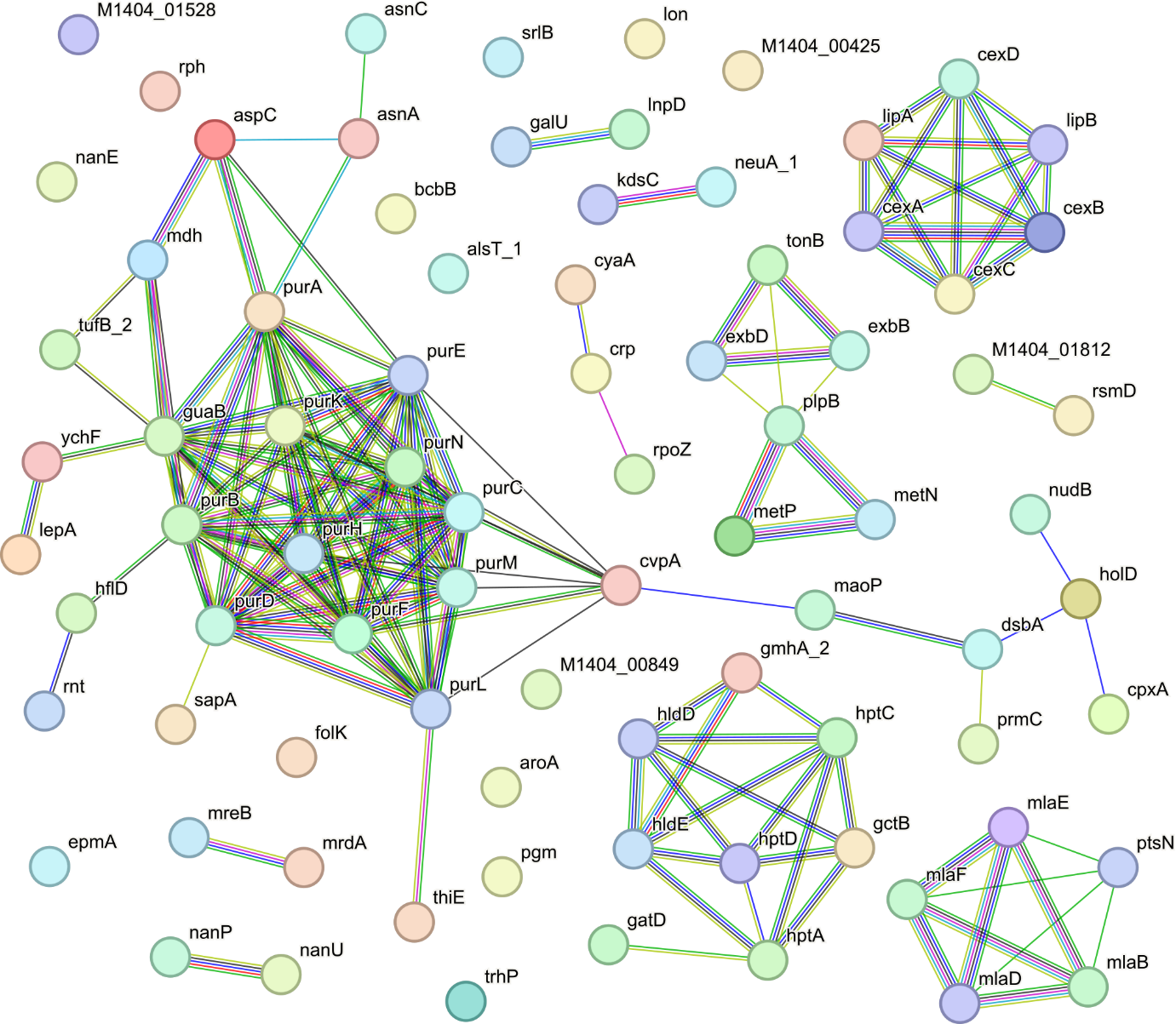

**S8 Fig.** STRING interaction map for *P. multocida* strain M1404 *in vivo* fitness genes. Genes were included if they were identified as important for M1404 survival in the bloodstream, liver, or spleen of BALB/c mice. The protein sequences of *in vivo* fitness genes were matched to *P. multocida* strain Pm70 in string, with 82 of the 93 proteins matching to a Pm70 protein. The lines between genes show the type of evidence for interaction; red line - fusion evidence, green line - neighbourhood evidence, blue line - co-occurrence evidence, purple line - experimental evidence, yellow line - text-mining evidence, light blue line - database evidence, black line - co-expression evidence.

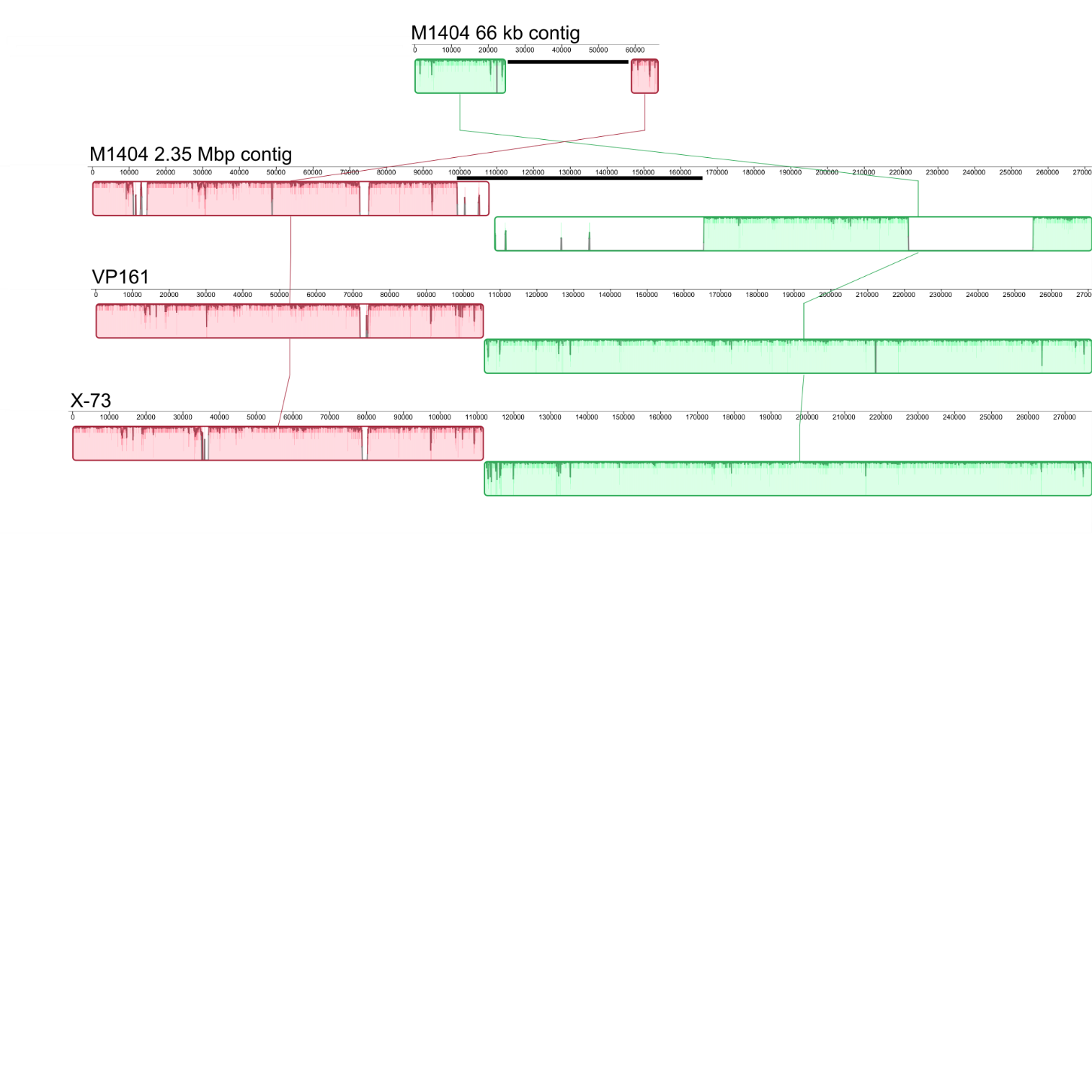

**S9 Fig.** Mauve alignment showing colinear parts of the 2.35 Mbp and 66 kb contigs generated in the *P. multocida* strain M1404 assembly with strains VP161 and X-73. The prophage regions are highlighted by black lines.
